# Fatty liver-mediated glycine restriction impairs glutathione synthesis and causes hypersensitization to acetaminophen

**DOI:** 10.1101/2023.01.16.524043

**Authors:** Alia Ghrayeb, Bella Agranovich, Daniel Peled, Alexandra C. Finney, Ifat Abramovich, Jonatan Fernandez Garcia, James Traylor, Shani Drucker, Sara Isabelle Fernandes, Natan Weissman, Y. Eugene Chen, Oren Rom, Inbal Mor, Eyal Gottlieb

## Abstract

Non-alcoholic fatty liver disease (NAFLD) affects nearly one third of the population worldwide. Understanding metabolic pathways involved can provide insights into disease progression. Untargeted metabolomics of livers from mice with early-stage steatosis indicated a decrease in methylated metabolites suggesting altered one carbon metabolism. The levels of glycine, a central component of one carbon metabolism, were lower in steatotic mice, in line with clinical evidence. Isotope tracing studies demonstrated that increased synthesis of serine from glycine is the underlying cause for glycine limitation in fatty livers. Consequently, the low glycine availability in steatotic livers impaired glutathione (GSH) synthesis under oxidative stress induced by acetaminophen (APAP), enhancing hepatic toxicity. Glycine supplementation mitigated acute liver damage and overall toxicity caused by APAP in fatty livers by supporting *de novo* GSH synthesis. Thus, early metabolic changes in NAFLD that lead to glycine depletion sensitize mice to xenobiotic toxicity even at a reversible stage of NAFLD.

## Introduction

Non-alcoholic fatty liver disease (NAFLD) is a leading cause of liver disease worldwide with numbers rising continuously (Younossi et al., 2018). While simple steatosis is reversible, NAFLD can progress to the more severe nonalcoholic steatohepatitis (NASH) characterized by hepatocyte damage and inflammation which leads to fibrosis that can eventually deteriorate to cirrhosis and/or hepatocarcinoma (Friedman et al., 2018). Indeed, NAFLD is becoming the leading cause of end-stage liver disease (Younossi et al., 2018) and there is a growing interest in the disease pathogenesis, its mechanisms and potential therapeutic strategies.

Metabolic changes are key to the development and exacerbation of NAFLD. One of the early hallmarks of fatty liver is the accumulation of lipid droplets in hepatocytes, giving rise to the disease’s name (Friedman et al., 2018). In addition to changes in fatty acid and carbohydrate metabolic pathways, which have been classically linked to NAFLD, amino acid metabolism has also emerged as a new feature in its pathogenesis (Gaggini et al., 2018; Hasegawa et al., 2020). Consistent changes in levels of specific amino acids in the circulation and in the liver were reported in both patients with NAFLD and pre-clinical models (Hasegawa et al., 2020). For example, there is considerable epidemiological evidence linking lower levels of circulating glycine to NAFLD, cardiometabolic diseases and metabolic syndrome (Gaggini et al., 2018; Li et al., 2018; Rom et al., 2022; Wittemans et al., 2019). Glycine based therapy has been shown to improve metabolic parameters in humans and pre-clinical models (Cruz et al., 2008; Rom et al., 2020; Takashima et al., 2016). However, the mechanism behind the characteristic decrease in glycine levels associated with NAFLD remains poorly understood.

The liver is the main site of drug metabolism and another important consequence of the metabolic changes under NAFLD is an effect on drug toxicity (Albadry et al., 2022; Massart et al., 2017; Naik et al., 2013). Since early stage NAFLD does not have clear clinical symptoms and is usually undiagnosed, the metabolic and functional changes that occur in the liver during early, reversible, stages of NAFLD, may have an effect on drug hepatotoxicity. In the current study, combining models of early-stage NAFLD and drug hepatotoxicity with stable isotope tracing studies, we elucidated the underlying metabolic mechanisms by which circulating and hepatic glycine is reduced in NAFLD, and examined the consequences of limited glycine availability on sensitivity to acetaminophen (APAP), a common analgesic and anti-pyretic drug, which is a leading cause of acute liver failure (Garcia-Roman and Frances, 2020; Larson et al., 2005).

## Results

### One carbon metabolism deficiency in fatty liver

To understand the initial undelaying metabolic changes that occur in NAFLD, we analyzed the liver metabolome at an early stage of the disease where only hepatic steatosis is present. To this end, mice were fed a high-fat high-sucrose diet, (“Western diet” - WD) or regular chow diet (CD) for 10 weeks. Mice fed the WD displayed increased body weight gain, elevated liver to body weight ratio, elevated levels of circulating alanine transferase (ALT), aspartate aminotransferase (AST), lactate dehydrogenase (LDH) and cholesterol (Figure 1A-F). Gross morphological analysis showed an increase in liver size and yellow coloration of the liver, indicating higher fat content, and histological analysis demonstrated simple steatosis as apparent by micro- and macro-steatosis as well as increased staining of neutral lipids (Figure 1G-J). Untargeted metabolic profiling of liver and plasma samples identified 350 compounds in the liver and 270 in the plasma. Of these, 112 metabolites in the liver and 90 metabolites in the plasma showed statistically significant changes in mice with WD-induced hepatic steatosis compared to control CD fed mice, revealing distinct metabolic profiles in each group (Figure 2A and 2B). Many methylated compounds were significantly lower in fatty livers (Figure 2C and 2D and TableS1 and S2; indicated in red). Pathway enrichment analysis of significantly altered metabolites in the liver and in the plasma revealed that serine and glycine metabolism as well as the methylation cycle (termed Methionine metabolism and Betaine metabolism) were highly ranked among the pathways identified (Figure 2E and 2F).

**Figure 1:**
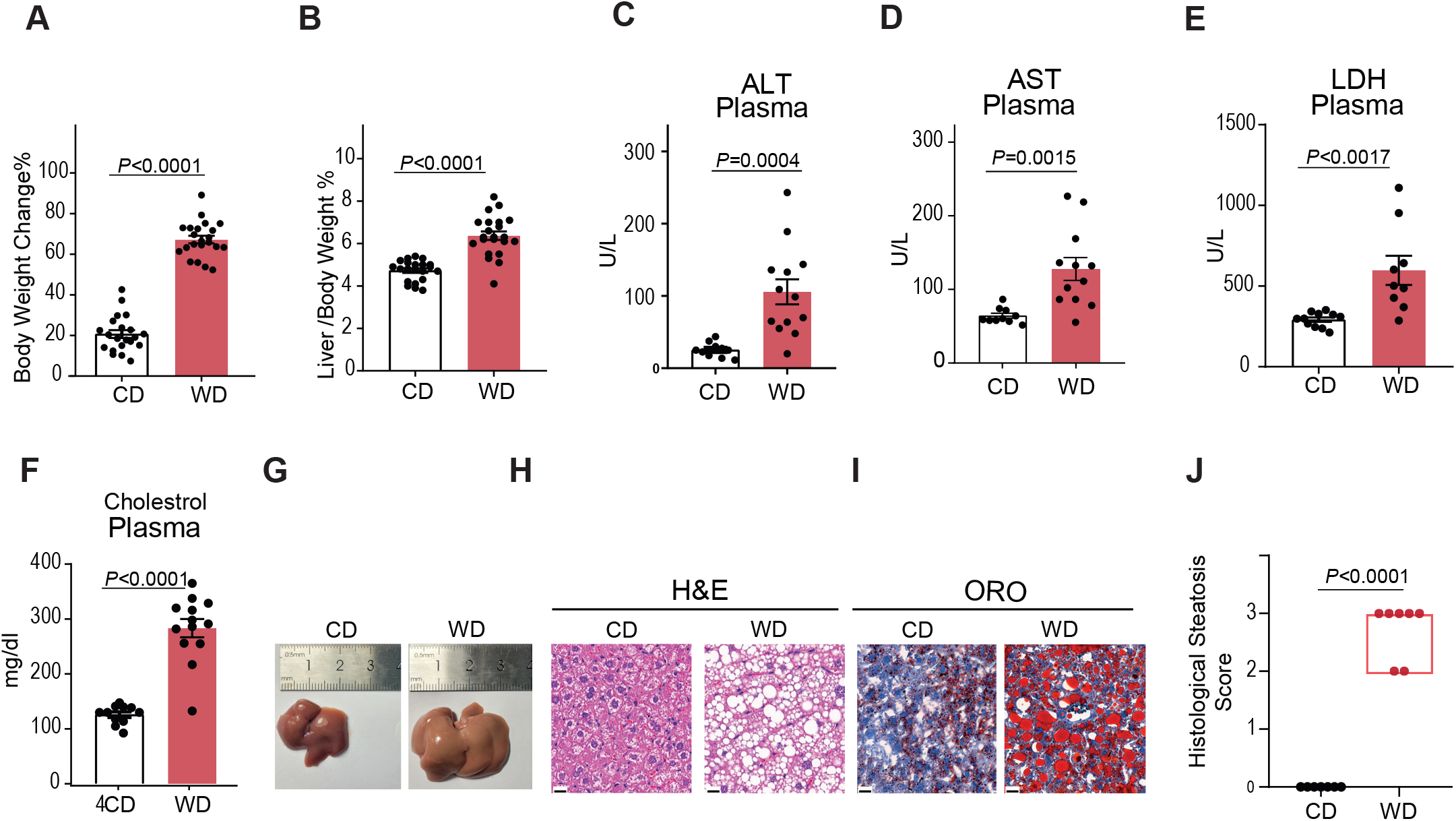
Simple steatosis in WD fed mice. **C57BL/6 mice were fed standard chow diet (CD) or western diet (WD) for 10 weeks. A and B,** Body weight change **(A)** and liver to body weight ratio **(B)** in CD compared to WD fed mice (n=21 and 22 respectively). **C-E,** Liver enzymes measured in the plasma, ALT (n=11 in CD and n=13 in WD group) **(C)**, AST (n=9 in CD and n=12 in WD group) **(D)** and LDH levels (n= 11 in CD and n=10 in WD group) **(E)**. **F,** Plasma cholesterol levels (n=11 in CD and n=13 in WD group). **G-J**, Gross liver morphology **(G)**, H&E **(H)**, ORO imaging (scale bars: 20 μm) **(I)** and Histological score of liver steatosis from representative CD or WD fed mice **(J)** (n=7 per group). Data presented as mean ± SEM, each point represents an individual mouse and P values were determined by two-tailed Student’s *t*-test for **(A-F).** Histological score is presented as median, each point represents an individual mouse, *P* values were determined by Mann–Whitney non-parametric test **(J)**.

**Figure 2:**
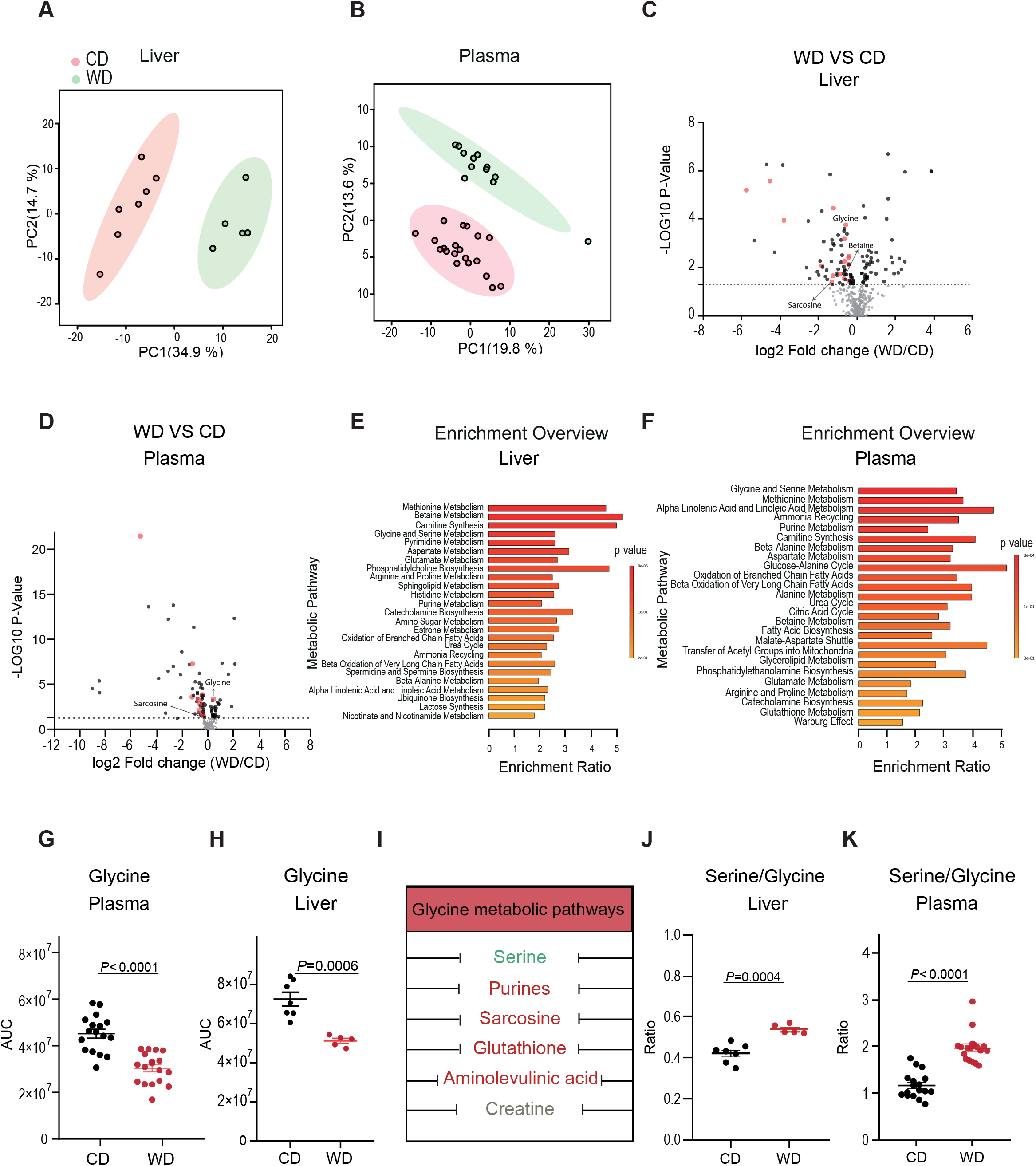
Downregulation of one carbon metabolism in mice with liver steatosis. Metabolic analysis of CD and WD fed mice. **A and B,** PCA of 350 chromatographic feature intensities in the liver (n=7 in CD and n=5 in WD group) **(A)** and 270 in the plasma (n=20 in CD and n=12 in WD group) **(B)**. **C-D,** Comparison of detected metabolites in CD vs WD fed mice in the liver (350 metabolites; n=7 in CD and n=5 in WD group) **(C)** and plasma (270 metabolites; n=20 in CD and n=13 in WD group) **(D)**. Scattered points represent metabolites, the x-axis indicates the log2 fold change of WD vs CD fed mice, the y-axis is the -log10 p-value, grey dots are metabolites with insignificant differences, black dots are metabolites with significant differences (-log (p-value) >1.3) and red dots are methylated metabolites with significant differences between CD and WD fed mice. **E-F,** Enrichment analysis based on metabolites that displayed significant differences in WD compared to CD fed group in the liver **(E)** and plasma **(F)**. For each SMPDB pathway, the bars show the enrichment ratio of the pathway in our dataset. **G,** Circulating levels of glycine measured using LC-MS analysis (n=17 in WT and n=18 in WD group). **H,** Hepatic levels of glycine (n=7 in CD and n=5 in WD group). **I,** Major metabolic outcomes of glycine (metabolites in red are decreased; green increased and grey are unchanged). **J,K,** Ratio of serine to glycine in the liver (n=7 in CD and n=5 in WD group) **(J)** and plasma (n=17 in CD and n=18 mice in WD group) **(K)**. Data presented as mean ± SEM. Each point represents an individual mouse. P values were determined by two-tailed Student’s t-test.

One of the main factors regulating methylation processes is availability of one carbon units which can be transferred to different substrates from folate and other methyl carriers (e.g. trimethylglycin/betaine; (da Silva et al., 2020; Ducker and Rabinowitz, 2017).The liver is known as a main site for one carbon metabolism, and alterations in lipid methylation have been reported in NAFLD (Alonso et al., 2017; da Silva et al., 2020; Walker, 2017). The interconversion of serine and glycine concomitantly with the transfer of a methyl group to/from tetrahydrofolate are key regulatory reactions in one carbon metabolism catalyzed by serine hydroxymethyltransferase (SHMT; these reactions occur both in the cytoplasm and in mitochondria by SHMT1 and SHMT2, respectively). Additionally, in the liver, glycine serves as a methyl donor through the glycine cleavage system which generates methyl tetrahydrofolate and CO_2_ (Ducker and Rabinowitz, 2017). Furthermore, glycine also serves as a methyl acceptor for glycine N-methyl transferase (GNMT) which produces monomethylglycine (sarcosine) and GNMT ablation in mice leads to hepatosteatosis (Rome and Hughey, 2022). Indeed, glycine sarcosine and betaine were among the significantly decreased metabolites in fatty liver compared to control (Figure 2C and 2D; Tables S1 and S2) and additionally to previously demonstrated results (Rom et al., 2020), glycine levels were significantly decreased in plasma of WD fed mice, even at this early stage of NAFLD (Fig. 2g, h).

### Glycine depletion in fatty liver is caused by an increase in serine synthesis from glycine

In an attempt to uncover the underlying mechanisms behind the characteristic decrease in glycine in NAFLD, we focused our attention on downstream products of glycine (Figure 2I). A consistent decrease in most glycine-derived metabolites was observed in WD fed mice (Figure 2I, Figure S1A-I) and is likely a consequence of limited glycine availability. In contrast, hepatic serine levels did not decrease while plasma serine levels were elevated, resulting in a high serine to glycine ratio (Figure 2J and 2K and Figure S1J and S1K). This indicates enhanced reverse SHMT activity as a potential underlying mechanism for the observed glycine depletion in the liver and plasma in NAFLD. To test this, stable isotope tracing using uniformly labeled ^13^C_2_ glycine was performed *in vivo* (Figure 3A). Glycine administration elevated total plasma glycine levels both in CD and WD fed mice and consequently liver glycine was also elevated albeit to a lower extent in WD fed mice (Figure 3B and 3C). Serine labeling in the liver indicated that a considerable fraction was synthesized from glycine (Figure 3D). Interestingly, an appreciable fraction of serine labeling was on all three carbons (M+3) indicative of combined glycine utilization via SHMT and the glycine cleavage system for *de novo* synthesis of serine (Figure 3D-F). Serine levels and labeling pattern in the plasma suggested a higher rate of serine synthesis from glycine in WD fed mice where two glycine molecules are used to synthesize one serine (Figure 3G). Analysis of glycine contribution to other downstream metabolites did not show an increase in the WD group (Figure S2). To test whether serine synthesis from glycine is the cause for the decreased glycine in NAFLD, we next inhibited this reaction using the pan-SHMT inhibitor, SHIN1 (Ducker et al., 2017). As expected, treating WD fed mice with SHIN1 blocked serine synthesis from glycine and significantly lowered steady state levels of serine in the liver and plasma (Figure 3H and 3I). Importantly, SHIN1 significantly elevated glycine levels in the liver and plasma of WD fed mice (Figure 3J and 3K), confirming our hypothesis on the etiology of glycine depletion in NAFLD due to enhanced serine synthesis.

**Figure 3:**
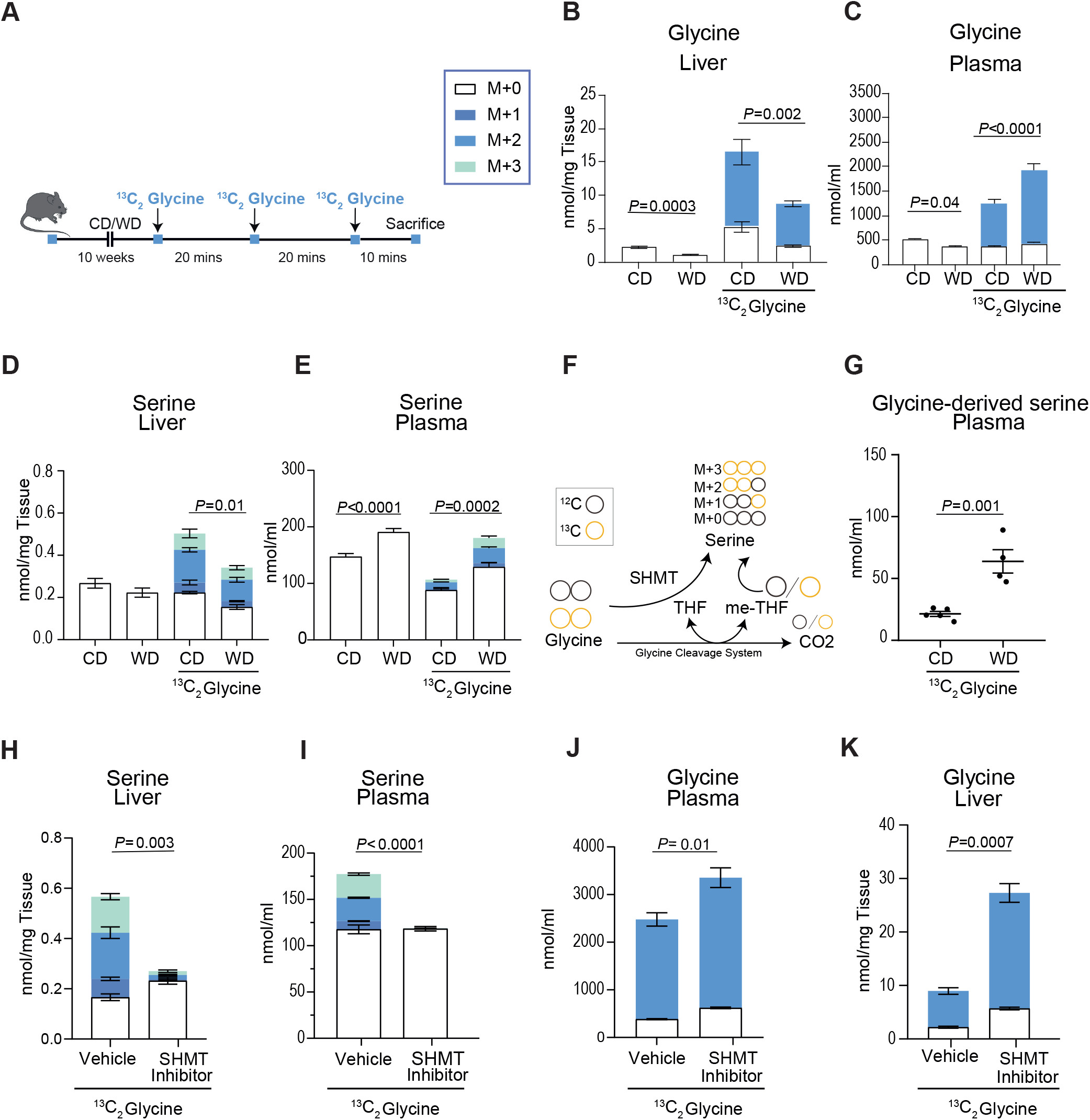
Increased serine synthesis causes glycine depletion in fatty liver. ^13^C_2_ glycine tracing in CD and WD fed mice. **A,** Schematic representation of the experimental design. **B-E,** Quantification and ^13^C isotope enrichment of glycine and serine in the liver (n=6 in CD and n=5 in all other groups) **(B and D)** and plasma (n=20 in CD, n=12 in WD, n=5 in CD-glycine and n=4 in WD-glycine group) **(C and E)**. **F,** Schematic representation of the possible number of ^13^C_2_-glycine-derived carbons in each serine isotopologue. **G,** levels of glycine-derived serine measured as the sum of weighted serine isotopologues as follow: ((M+1) nmol/ml _X_ 1) + ((M+2) nmol/ml _X_ 1) + ((M+3) nmol/ml _X_ 2). Each point represents an individual mouse. **H-K,** Metabolic analysis of the liver and plasma 70 minutes following SHMT inhibitor administration (n=3 in vehicle and n=4 in SHMT inhibitor group). Data presented as mean ± SEM. *P* values determined by two-tailed Student’s *t*-test **(B, G-K)** or one-way analysis of variance (ANOVA), followed by Tukey’s test **(D and E)**.

### Glycine limitation in steatotic hepatocytes leads to susceptibility to xenobiotic mediated oxidative stress

An important consequence of glycine deprivation is the inhibition of GSH synthesis (Mardinoglu et al., 2017; Rom et al., 2022; Rom et al., 2020; Sekhar et al., 2011). Synthesized from the three amino acids – glutamate, cysteine and glycine, GSH, is the cell’s major antioxidant (Figure S3A). Indeed, hepatic GSH was decreased concomitantly with glycine, but not with glutamate or cysteine in WD fed mice (Figure S1C, Figure S3B and S3C). To examine the biochemical consequences of lipid accumulation on glycine and GSH metabolism in hepatocytes, we used the immortalized mouse hepatocyte cell line, AML12. Cells incubated with increasing concentrations of palmitate for 24 hrs demonstrated elevated levels of triglycerides and formed lipid droplets, characteristics of fatty liver (Figure 4A and 4B). In line with our *in vivo* findings, intracellular levels of glycine and GSH were significantly decreased in palmitate-loaded AML12 cells (Figure 4C-E). This was associated with increased oxidative stress, as indicated by enhanced fluorescence of dihydroethidium (DHE), a superoxide probe (Figure 4F). Hepatic GSH is a key player in xenobiotic neutralization and a decrease in GSH levels could have deleterious consequences on drug metabolism and their toxicity (Lushchak, 2012; Wu et al., 2004). Acetaminophen (APAP) is a widely used analgesic and anti-pyretic drug. It is commonly prescribed and considered to be safe, nevertheless, APAP overdose is the leading cause of acute liver failure worldwide (Garcia-Roman and Frances, 2020; Larson et al., 2005). In safe doses, APAP is converted in the liver to non-toxic compounds while a small fraction of the drug is metabolized to a reactive free radical, N-acetyl-p-benzoquinone imine (NAPQI), which is then neutralized by GSH (Figure 4G). However, due to APAP overdose, the high levels of NAPQI exceed GSH availability causing oxidative damage to the liver. Indeed, APAP administration enhanced superoxide formation in AML12 cells, which increased when APAP was co-administered with palmitate (Figure 4H). The combined palmitate and APAP administration was also significantly more toxic than each agent alone (Figure 4I).

**Figure 4:**
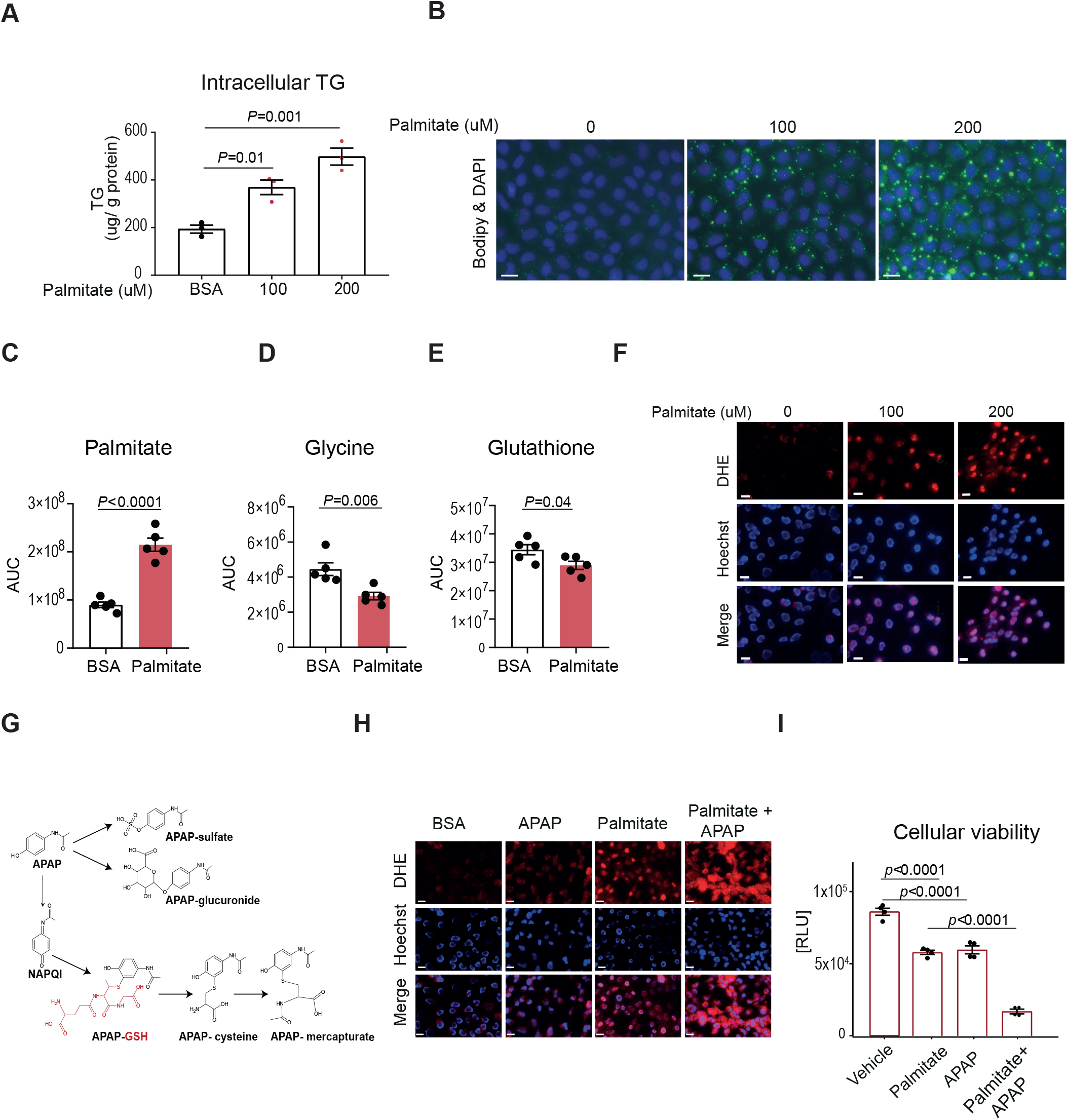
Lipid loading decreases intracellular levels of glycine and GSH in cultured hepatocytes and increases sensitivity to APAP. AML12 cells incubated with BSA or palmitate for 24 hrs. **A,** Cellular triglyceride content. **B,** Visualization of intracellular neutral lipids using BODIPY 493/503 staining (scale bar: 20 μm). **C-E,** LC-MS analysis of the indicated metabolites. **F,** Superoxide was detected with dihydroethidium (DHE, red) and nuclei stained with Hoechst (blue) in live cells following indicated treatments (scale bar: 20 μm). **G,** APAP metabolism in the liver. **H,** DHE (red) and Hoechst (blue) staining in live cells following indicated treatments (scale bar: 20 μm). **I,** Cell viability measured using CellTiter-Glo™ assay. All graphs and images are representative of three different experiments, at least three wells in each experiment. Data presented as mean ± SEM. *P* values were determined by ANOVA, followed by Tukey’s test **(A, I)** or two-tailed Student’s *t*-test **(C-E)**.

Epidemiological evidence suggests that NAFLD patients have an increased risk for APAP toxicity, yet results from pre-clinical studies are inconclusive (Garcia-Roman and Frances, 2020; Ito et al., 2006; Kim et al., 2017b; Michaut et al., 2014). We therefore examined weather WD fed mice, which developed simple steatosis and had lower hepatic GSH (Figure S1C), presented increased sensitivity to APAP. First, we determined APAP metabolism and toxicity in CD fed mice at increasing doses from 300 to 600 mg/kg. These doses are equivalent to 2-4 times the maximum single dose recommendations in humans (Nair and Jacob, 2016) (www.nhs.uk/medicines/paracetamol-for-adults/about-paracetamol-for-adults/). APAP and its main downstream metabolites following sulfate or glucuronide conjugation (Figure 4G) were detected in the plasma in a dose-dependent manner (Figure S4A-C). However, catabolic products of NAPQI conjugated to GSH (APAP-GSH), namely APAP-cysteine and APAP-meracapturate, reached their maximum levels already at 300 mg/kg (Figure S4D and 4E) while APAP-GSH itself was not detected in the plasma. Metabolism of APAP occurs mainly in cells proximal to the central vain (zone 3) that express the cytochrome p450 enzyme, CYP2E1 (Blazka et al., 1996; Hinson et al., 2010). Histological analysis at 24 hrs post APAP administration demonstrated cell death in zone 3 at all doses, however, the 450 and 600 mg/kg doses caused wider necrotic changes (ballooning degeneration) beyond zone 3 (Figure S4F). Furthermore, signs of distress were observed in APAP-treated mice receiving 450 and 600 mg/kg (not shown) while only one out of 7 CD fed mice treated with 300 mg/kg showed minor signs of distress (Supp movie 1A). Therefore, we chose an APAP dose of 300 mg/kg which was well-tolerated in CD fed mice.

To determine the effect of liver steatosis on APAP hepatotoxicity, mice fed with WD or CD for 10 weeks received a single dose of APAP (300 mg/kg) or saline (Figure 5A). APAP administration to WD fed mice resulted in extremely high values of circulating liver enzymes, ALT and LDH 24 hrs post injection, indicating considerable tissue damage (Figure 5B and 5C). Examination of the liver gross morphology revealed visible hemorrhages in livers from mice on WD, but not on CD (Figure 5D). Histological analysis demonstrated hemorrhages present in zone 3 with a significantly higher histological score in mice on WD compared to those on CD (Figure 5E and 5F). At that point, 7 out of 13 of the APAP treated WD fed mice showed visible clinical signs and reached endpoint (Supp movie 1B). Pharmacological studies revealed that GSH was depleted 3 hrs following APAP treatment in both CD and WD fed mice (Figure 5G). Nevertheless, and in line with the *in vitro* data, WD fed mice had a lower capacity to recover GSH levels and they demonstrated elevated redox stress as indicated by oxidized to reduced glutathione ratio (GSSG/GSH) (Figure 5G and 5H).

**Figure 5:**
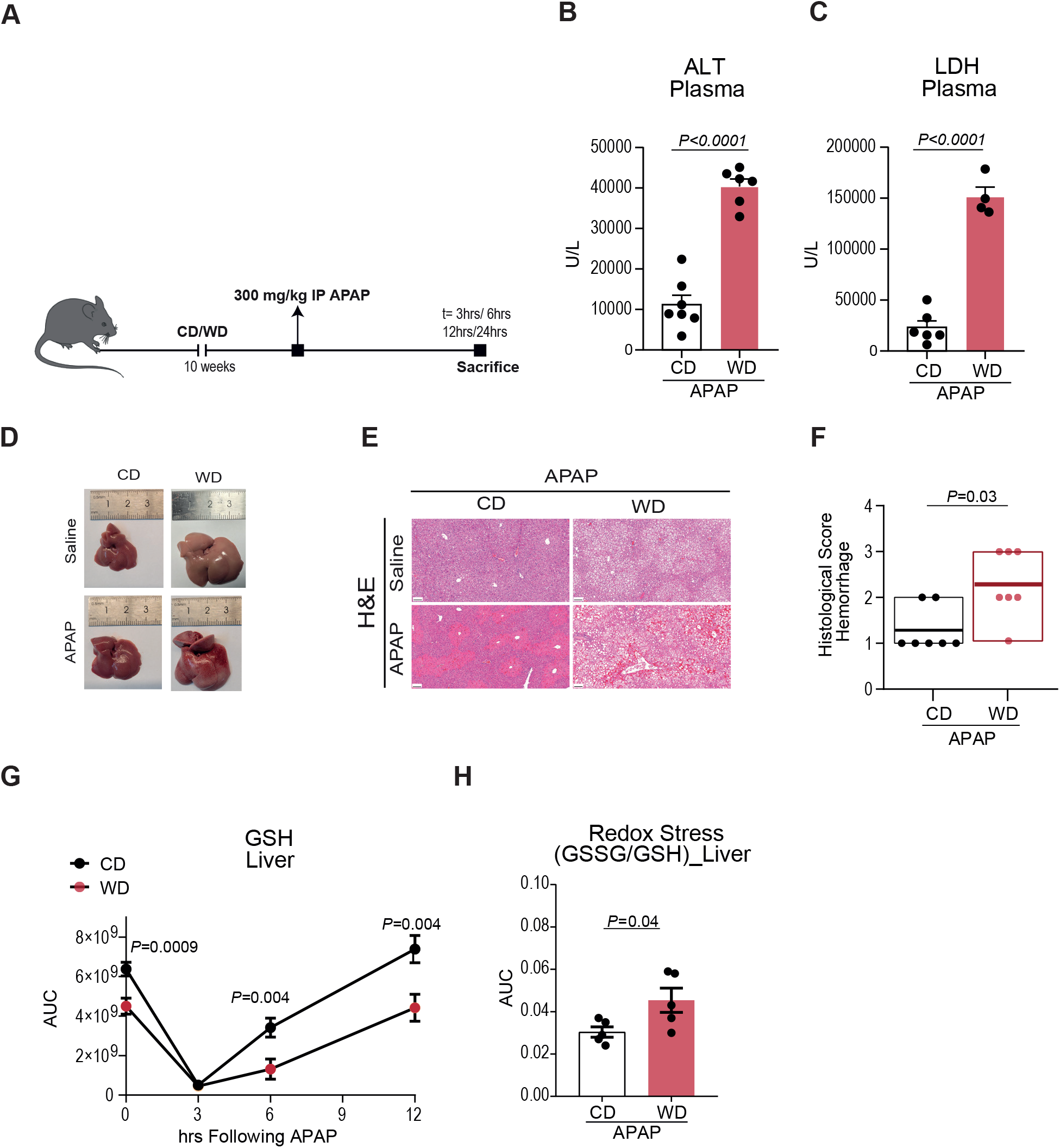
NAFLD mice display decreased GSH recovery, increased redox stress and hepatotoxicity. APAP administration (300 mg/kg) after 10 week of CD or WD. **A,** Schematic representation of the experimental design. **B and C,** Plasma levels of ALT (n=7 in CD and n=6 in WD group) **(B)**, and LDH (n=6 in CD and n=4 in WD group) **(C)**. **D-F,** Gross liver morphology **(d),** H&E staining (scale bars: 100 μm) **(E)**, and histological score of hemorrhage 24 hrs following APAP injection (n=7 mice per group) **(F). G,** Hepatic GSH at indicated time points (t=0 hr: n=12 in CD and n=7 in WD group; t= 3 hrs: n=13 in CD and n=7 in WD group; t=6 hrs n=11 in CD and n=7 in WD group; t=12 hrs: n=5 per group). **H,** Redox stress index following 12 hrs of APAP administration (n= 5 per group). Data presented as mean ± SEM, each point represents an individual mouse*, P* values were determined by two-tailed Student’s *t*-test for **(B, C, G and H).** Histological scores are presented as median **(F)**, each point represents an individual mouse and *P* values were determined by Mann–Whitney non-parametric test.

### Glycine administration rescues hepatic GSH levels and mitigates APAP hepatotoxicity

To determine whether lower glycine availability in steatotic livers is the underlying mechanism for the increased APAP sensitivity, steatotic mice were treated with 1 g/kg glycine (Caldow et al., 2016; Rom et al., 2022; Rom et al., 2020) immediately after APAP administration and 3 hrs later, when GSH levels are dramatically depleted and its recovery is initiated (Figure 5G and Figure 6A). Oral glycine administration significantly elevated hepatic glycine and doubled the recovery rate of GSH 6 hrs post APAP administration (Figure 6B and 6C). This alleviated APAP-induced redox stress as indicated by GSSG/GSH ratio and protected from lipid oxidation in the liver as measured by GSH conjugated to the lipid peroxide product 4-hydroxynonenal (4-HNE) (Figure 6D and 6E). These results indicate that the limitation on GSH synthesis by low glycine availability in fatty liver was alleviated by exogenous glycine. Functionally, glycine administration dramatically protected from liver damage as measured by circulating ALT and LDH and from APAP-induced liver hemorrhage and necrosis (Figure 6F-K). Consequently, glycine treatment improved clinical signs and only 2 out of 15 WD fed, APAP-treated mice reached endpoint, improving the survival rate from 46% to 87% (Figure 6L and Supp movie. 1C).

**Figure 6:**
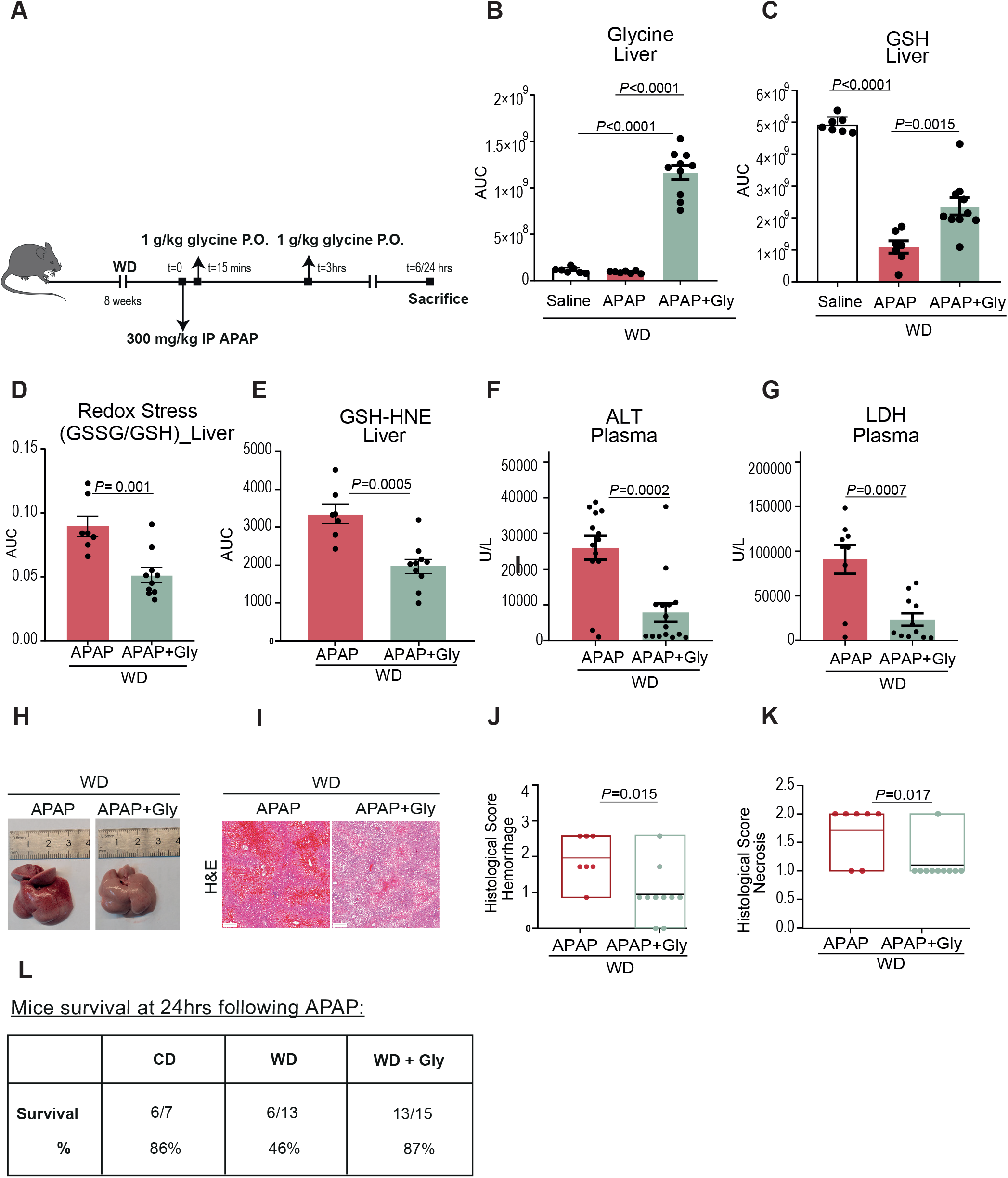
Glycine administration rescues hepatic GSH levels and mitigates APAP hepatotoxicity. WD fed mice gavaged with 1g/kg glycine following APAP injection. **A,** Schematic representation of experimental design. **B and C,** Metabolic analysis of livers 6 hrs following administration of APAP (n=7 in Saline, n=7 in APAP and n=10 in APAP+Gly group). **D and E,** Redox stress index and 4-HNE conjugated to GSH (n=7 in APAP and n=10 in APAP+Gly group). **F and G,** Liver enzymes in the plasma 24 hrs following administration of APAP and glycine, ALT (n=13 in APAP and n=15 in APAP+Gly group) **(F),** and LDH (n=9 in APAP and n=11 in APAP+Gly group) **(G). H-K,** Liver morphology **(H)**, H&E from representative mice (scale bar: 200um) **(I),** and histological score of hemorrhage **(J)** and necrosis **(K)** 24 hrs following APAP and glycine administration (n=7 in APAP and n=10 in APAP+Gly group). **L,** Survival rate at 24 hrs following APAP injection. Data presented as mean ± SEM, each point represents an individual mouse, values were determined by ANOVA followed by Tukey’s test **(B and C)** or two-tailed Student’s *t*-test **(D-G)**. Histological scores are presented as median, each point represents an individual mouse, *P* values were determined by Mann–Whitney non-parametric test **(J and K)**.

## Discussion

We studied the metabolic profile of mice at the early stages of NAFLD, when only simple steatosis has developed. While this can be a stable and even reversible stage, it is the first towards a more advanced liver disease (Friedman et al., 2018). Metabolic changes have been extensively studied and naturally have mainly focused on central pathways that would result in lipid accumulation in hepatocytes, namely carbohydrate and fatty acid metabolism. Expectedly, we detected changes in various dietary fatty acids and acyl-carnitines (Figure 2 and Table S1). Analysis of the liver metabolome demonstrated a decrease in methylated metabolite species in livers from WD fed mice which indicated changes in one carbon metabolism. The liver is a main site of one carbon metabolism and this pathway, which is also connected to phospholipid synthesis, has been implicated in NAFLD pathophysiology (da Silva et al., 2020). Serine and glycine are major players in the transfer of one carbon units (Ducker and Rabinowitz, 2017). Several studies of circulating metabolites have identified lower glycine as a predictive biomarker NASH and fibrosis in patients (Gaggini et al., 2018; Hasegawa et al., 2020; Zhou et al., 2016b). Our results show that the decrease in circulating glycine occurs early in NAFLD, already at the stage of simple steatosis. Previously, we and others have demonstrated the beneficial effect of glycine supplementation in NASH, improving liver clinical parameters (Rom et al., 2020; Takashima et al., 2016; Zhou et al., 2016a). Here we provided an unrecognized metabolic mechanism that explains the cause for the specific decrease in glycine in NAFLD.

It has been previously suggested that the decrease in circulating glycine in cardiometabolic disease is secondary to the elevation in branched chain amino acids (BCAAs) which is commonly reported in obesity and type 2 diabetes (White et al., 2020). Yet, in our fatty liver model, we did not detect a consistent increase in circulating BCAAs, indicating that in hepatic steatosis, changes in glycine metabolism could occur independently of BCAAs metabolism.

Our stable isotope tracing studies revealed that glycine is a substantial source for serine synthesis in fatty liver. These studies, further indicate that glycine is also consumed through the glycine cleavage system, meaning that two glycine molecules are utilized to produce one serine with one carbon released as CO_2_. In our studies, circulating serine was significantly elevated and labeled considerably with carbons originating from glycine. Moreover, the inhibition of serine synthesis was sufficient to elevate glycine levels. Therefore, the metabolic flux of glycine to serine, mediated by reverse SHMT activity and glycine decarboxylase is elevated in fatty liver disease and is the major cause for the characteristic decrease in circulating glycine.

GSH levels are known to decrease in NASH but were also demonstrated to decrease under simple steatosis induced by choline deficient diet which does not involve inflammation and fibrosis (Grattagliano et al., 2008). Here we show that the decrease in GSH also occurs when steatosis is induced by a more clinically relevant Western-type diet, demonstrating it as a characteristic trait of early stages of NAFLD. GSH is essential for xenobiotic detoxification in the liver (Lushchak, 2012; Wu et al., 2004) and human epidemiological studies correlated fatty liver disease with increased sensitivity to xenobiotic (Massart et al., 2017; Merrell and Cherrington, 2011). Cysteine is considered to be rate limiting in GSH synthesis and its administration in the form of N-Acetyl-Cysteine (NAC), is the standard of care for APAP overdose (Garcia-Roman and Frances, 2020). However, of the three amino acids required to produce GSH, only glycine is lower in fatty liver. Interestingly, glycine can also support *de novo* GSH production by enabling cysteine synthesis from glycine-derived-serine and homocysteine via the transsulfuration pathway (Figure S5A) (Ducker and Rabinowitz, 2017). Indeed, supplementing steatotic mice with glycine led to an increased levels of liver cystathionine and cysteine (Figure S5B and S5C). Furthermore, glycine administration decreased metabolic markers of oxidative stress and improved the clinical outcome of APAP toxicity (Figure 5). This suggests that monitoring circulating glycine should be taken into consideration when administrating analgesics to patients with NAFLD even at early stages of simple steatosis.

To summarize, our study mechanistically linked glycine limitation in NAFLD with GSH synthesis and xenobiotic sensitivity. We demonstrated that supplementing glycine to WD fed mice restored circulating and hepatic glycine levels, increased recovery of GSH following APAP administration and protected steatotic mice from APAP toxicity.

## Acknowledgments

This study was supported by grants from the Israel Science Foundation (ISF) (#824/19 to E.G.), the Michigan-Israel Partnership for Research and Education (E.G., I.M., O.R., Y.E.C.), the Laura and Isaac Perlmutter Foundation (E.G.), the Israel Council for Higher Education (VATAT to A.G.) and the National Institute of Health (R00 HL 150233 and R01 DK134011 to O.R.). We thank the Biomedical Core Facility (BCF) and the preclinical facility at the Ruth and Bruce Rappaport Faculty of Medicine, Technion – Israel Institute of Technology for their services.

## Author contributions

A.G. designed the study, preformed most experiments, collected and analyzed the data, and co-wrote the manuscript with other authors. D.P., A.F. carried out *in vitro* experiments, collected data and analyzed imaging experiments. I.A., B.A. supervised LC-MS runs and developed the analytical methods. J.T. scored the histological slides. S.F., N.W., S.D. assisted with *in vivo* experiments. J.G. assisted with metabolomics dataset analysis. O.R., Y.E.C discussed the data and commented on the manuscript. I.M., E.G. supervised the project and funding acquisition, designed studies, and wrote the manuscript.

## STAR METHODS

### Animal studies

8-week-old C57BL/6JOlaHsd male mice were purchased from Envigo. All animal procedures were performed in accordance with guidelines established by the NIH on the care and use of animals in research, as confirmed by the Technion Institutional Animal Care and Use Committee.

To induce fatty liver disease, mice (n=3 per cage) were fed ad libitum either standard chow diet (altromin, 1320) or western diet (Envigo, TD.88137) for 10 weeks. For glycine tracing experiments, mice were injected intraperitoneally thrice with 100 mg/kg ^13^C_2_ glycine (Sigma-Aldrich, 283827) in saline (0.9% w/v of NaCl in water) at 20 minutes intervals and sacrificed 10 minutes following the third injection. For SHMT inhibitor experiments, mice were injected intraperitoneally with 100 mg/kg SHIN1 (MedChemExpress, HY-112066A), 20 minutes prior to ^13^C_2_ glycine administration.

For APAP experiments, food was removed from the cages 16 hrs before administration. On the day, 15 mg/kg APAP (Sigma-Aldrich, A7085) was dissolved in saline and kept at 37°C until administrated intraperitonealy at 300 mg/kg dose. Saline was used as control. Food was returned to cages 2 hrs follwoing APAP administration. For glycine treatment, 1 g/kg glycine (Sigma-Aldrich, G7126) was administered by oral gavage 15 minutes and 3 hrs following APAP injection. Mice were sacrificed at indicated time points.

### Histology and immunohistochemistry

The livers were dissected, and the middle lobe fixed in 4% paraformaldehyde (PFA) for 48 hrs, paraffin embedded and sectioned at 5 μm thickness. Mounted tissue sections were stained with hematoxylin and eosin (H&E) for histological analysis. For Oil Red O (ORO) staining, PFA-fixed liver samples were cryoprotected in 20% sucrose at 4°C overnight, liquid nitrogen snap-frozen in Tissue Freezing Medium (Leica, 14020108926) and stored at −80°C until ready for cryosectioning at 10 μm thickness. Mounted tissue sections were stained with ORO (Sigma-Aldrich, 01391) as described before (Ma et al., 2019). Slides were nuclear counterstained in Harris Hematoxylin (Sigma-Aldrich, MHS1) and mounted in aqueous mounting media (Leica, 94-9990402).

### Liver histology scoring

Liver histology was scored in H&E-stained, paraffin-embedded sections. Steatosis was scored from 0-3, APAP induced liver injury (AILI) from 1-5 and hemorrhage from 1-3, according to previously published papers (Bedossa et al., 2012) (Latchoumycandane et al., 2007) (Hoque et al., 2012). Pathologists were blinded to the experimental groups.

### Serum analysis

Blood samples were collected in serum collection tubes with clotting activator (grenier, 450533). Tubes were left at room temperature for 20 minutes to allow for coagulation. To separate the serum from blood clot, tubes were centrifuged at 2000g for 15 minutes at 4 °C. Liver enzyme and cholesterol analysis were performed by clinical biochemistry lab at Rambam Health Care Campus. In hemolytic blood samples, the LDH and AST levels were excluded from the analysis to avoid false positive effect (which explains the variability in the number of mice (n) per group between the graphs of liver enzymes in Fig. 1, Fig. 5 and Fig. 6).

### Sample preparation and metabolite extraction

For tissue collection, mice were sacrificed via cervical dislocation and liver tissues were immediately collected, gallbladders were removed, livers were then rinsed with cold PBS, cut into different lobes, snap-frozen in liquid nitrogen and kept in −80□°C. To maximize accuracy and avoid variability, metabolomic analysis was performed only on the left lobe of the liver. Approximately 40 mg (±SD of 1 mg) of frozen tissue was added to CK14 homogenizing tubes (Sarstedt Inc, 72.694.305) containing 1.4 mm ceramic beads (Bertin Corp, P000926-LYSK0-A) which were prefilled with 1000ul of cold (−20°C) metabolites extraction solvent of methanol: acetonitrile: water at 5:3:2 ratio. Samples were homogenized at 4°C in a Precellys 24 tissue homogenizer (Bertin Corp), homogenization conditions were set to 3 cycles, 20 seconds each, 7200 rpm with a 60 second gap between each of the three cycles to preserve low temperature. Homogenates were centrifuged at 18,000 g for 15 min at 4°C. The supernatant was collected in a microcentrifuge tube and centrifuged again. The cleared supernatants were transferred to glass HPLC vials and kept at −80°C until LC-MS analysis. Plasma was collected in collection tubes with heparin lithium (BD, 450535) and then centrifuged at 2000g for 15 minutes at 4°C.

For metabolic extraction, plasma was diluted 1:10 in cold extraction solvent of methanol: acetonitrile at 75:25 ratio, vortexed for 10 minutes and immediately centrifuged at 18,000*g* for 10 min at 4□°C. The supernatant was collected for LC–MS analysis.

### Metabolites quantification

For glycine and serine quantification in the liver, calibration standards *of* ^13^C_2_ glycine (Sigma-Aldrich, 283827) and 13C_3_ serine (Cambridge Isotope Laboratories, Inc, CLM-1574-H) were prepared in liver tissue homogenates as matrix, at a concentration range of *0.58-1200uM*. Calibration curves were prepared in homogenates from both healthy liver and fatty liver to overcome possible matrix effect. *For glycine and serine quantification in the plasma*, calibration standards were prepared in plasma matrix (CD fed mice) at 0.38-1200uM concentration range. *Samples were processed identically to liver samples*.

### Targeted metabolomics

For polar metabolites detection, LC-MS metabolomics analysis was performed as described previously (Mackay et al., 2015). ThermoFisher Scientific Ultimate 3000 high-performance liquid chromatography (HPLC) system coupled to Q-Exactive Orbitrap Mass Spectrometer (ThermoFisher Scientific Fisher Scientific) was used with a resolution of 35,000 at 200 mass/charge ratio (*m/z*), electrospray ionization, and polarity switching mode to enable both positive and negative ions across a mass range of 67 to 1000 *m/z*. HPLC setup consisted ZIC-pHILIC column (SeQuant; 150 mm × 2.1 mm, 5 μm; Merck), with a ZIC-pHILIC guard column (SeQuant; 20 mm × 2.1 mm). 5 ul of biological extracts were injected and the compounds were separated with mobile phase gradient of 15 min, starting at 20% aqueous (20 mM ammonium carbonate adjusted to pH 9.2 with 0.1% of 25% ammonium hydroxide) and 80% organic (acetonitrile) and terminated with 20% acetonitrile. Flow rate and column temperature were maintained at 0.2 ml/min and 45°C, respectively, for a total run time of 27 min. All metabolites were detected using mass accuracy below 5 ppm. ThermoFisher Scientific Xcalibur was used for data acquisition.

For the detection of APAP-GSH peak, mice were injected with labeled [U-13C] glutamine (Cambridge Isotope Laboratories, Inc, CLM-1822-H) As glutamine is a carbon donor for GSH synthesis, a mass shift of 5 [Da] in APAP-GSH mass was used to validate the peak of APAP-GSH.

For cysteine detection- The mass spectrometer (Thermo Q-Exactive Orbitrap) was operated in a positive polarity mode by t-SIM (targeted-selected ion monitoring). The selected mass transitions were m/z 122.02709 for M+0, m/z 123.03038 for M+1, m/z 124.03374 for M+2 and m/z 125.03709 for M+3. HPLC setup consisted of ZIC-pHILIC column (SeQuant; 150 mm × 2.1 mm, 5 μm; Merck), Mobile phase A: 0.1% formic acid v/v in water. Mobile B: acetonitrile. The flow rate was kept at 200□μl/□min and gradient were as follows: 0 min 50% of B, 8□min 20% of B, 12 min 20% of B, 14 min 50% of B, 18-21□min 50% of B.

### Metabolic data analysis

TraceFinder 4.1 (Thermo Scientific) was used for analysis. Peak areas of metabolites were determined using the exact mass of singly charged ions. The retention time of metabolites was predetermined on the pHILIC column by analyzing an in-house mass spectrometry metabolite library consisting of commercially available standards. For data normalization, samples were normalized to mg tissue. Correction for natural abundance effect in isotope tracing experiment was done using Metabolite AutoPlotter 2.5 (Pietzke and Vazquez, 2020).

### Untargeted metabolomics

Untargeted metabolomics analysis was carried out using Compound Discoverer software 3.3 (Thermo Scientific). Retention times were aligned across all data files with a maximum shift of 2 min and mass tolerance of 5 ppm using the pool samples as a reference file. Unknown compound detection (minimum peak intensity of 1□×□10^5^) and grouping of compounds were performed across all samples with the following main settings (all other parameters were kept as default values): mass tolerance of 5 ppm, retention time tolerance of 0.2 min compound detection: M+H and M-H ions only, peak rating filter was set to 4. Missing values were filled using the software’s ‘Fill Gap’ feature (mass tolerance of 5 ppm and signal/noise tolerance of 1.5). 1081 and 1200 features were detected in the plasma and liver extracts respectively. Metabolites annotation was done as follows, with decreasing confidence level: First, by matching the mass and retention time of observed signal to an in-house library generated using commercial standards (mass tolerance of 5 ppm and retention time tolerance of 0.5 min) or by matching fragmentation spectra to mzCloud (www.mzcloud.org) with precursor and fragment mass tolerance of 10 ppm and match factor threshold of 80. Compounds with no fragmentation spectra data were annotated using Human Metabolom Database (https://hmdb.ca/), BioCyc (https://biocyc.org/) and Kyoto Encyclopedia of Genes and Genomes (KEGG https://www.genome.jp/kegg/) applying filtering of the software’s’ mzlogic score higher than 50 and with less than three possible candidates. After completing metabolites filtering and annotation process, 270 and 350 metabolites were annotated in the plasma and liver receptively.

### Cell culture

AML-12 cells were obtained from ATCC (CRL-2254) and cultured according to the manufacturer’s recommendations in DMEM: F12 media (Biological Industries, 01-170-1A) supplemented with with 10% fetal bovine serum (FBS) (ThermoFisher Scientific. 10270106), 100 U/mL penicillin (Biological Industries, 03-031-1B), 100 μg/mL streptomycin (Biological Industries, 03-031-1B), 40 ng/mL dexamethasone (Sigma-Aldrich, D4902), ITS-X (1:100, ThermoFisher Scientific, 41400045) and 2mM Glutamine (Biological Industries, 03-020-1A).

For metabolomic experiments, cells were seeded in 6 well plates at a density of 2 × 10^5^ cells per well. On the following day, growth medium was replaced with metabolic medium: EBSS (Biological Industries, 02-0101-1A), 1% FBS, 100 U/mL penicillin, 100 μg/mL streptomycin, non-essential amino acids (1:50, Biological Industries, 01-325-1B), vitamin solution (1:50, Biological Industries, 01-326-1B), 0.25 mM L-serine (Sigma-Aldrich, S4500), 0.25 mM glycine (Sigma-Aldrich, G7126), 1 mM glutamine (Biological Industries, 03-020-1A), 1 mM sodium pyruvate (Biological Industries, 03-042-1B), ITS-X (1:100, ThermoFisher Scientific, 41400045), 40 ng/mL dexamethasone, 1 μg/mL glutathione (Sigma-Aldrich, G6013), 0.3 ng/mL ammonium metavanadate (Sigma-Aldrich, 204846), 0.25 nM manganese chloride (Sigma-Aldrich, 244589), 2.5 mg/L ascorbic acid (Sigma-Aldrich, A4403). The following day medium was replaced with fresh metabolic medium supplemented with 200uM palmitate-BSA or BSA only for 24 hrs. Palmitate-BSA conjugation was performed as previously described (Kim et al., 2017a)). Metabolic extraction from cells using methanol: acetonitrile: water at 5:3:2 ratio and LC-MS analysis were performed as previously described (Mackay et al., 2015). For data normalization, raw data files were processed with Compound Discoverer 3.3 to obtain total compounds peak area for each sample. Each metabolite peak area value was normalized to total measurable ions in the sample.

### Lipid analysis

Cells were plated on 8 well chamber slides (ThermoFisher Scientific, 154526PK) at a density of 1.5 × 10^4^ cells per well. Following 24 hrs of incubation with palmitate-BSA or BSA only, cells were fixed with 4% PFA for 10 minutes, and stored at ***4***°C until analysis. Cellular lipids were stained with 5uM BODIPY 493/503 (ThermoFisher Scientific, D3922) according to manufacturer’s instructions and nuclei were stained with DAPI 5 ug/ml (Sigma-Aldrich, D9542).

### Superoxide analysis

AML12 cells were plated onto 8 well chamber slides (ThermoFisher Scientific, 154526PK) at density of 1.5 × 10^4^ cells per well. The next day, cell growth media was replaced with metabolic medium supplemented with either BSA, palmitate-BSA, 10 mM APAP or APAP and palmitate-BSA. Following 24hrs, cells were washed 3 times with warm HBSS media (ThermoFisher Scientific, 14025050) and incubated with 5uM Dihydroethidium (DHE) (Sigma-Aldrich, D7008) for 30 mins at 37 °C in the dark. Cells were then washed again with warm HBSS media, and live nuclei were stained with Hoechst (ThermoFisher Scientific, R37165). Cells were maintained in HBSS with Hoechst during image acquisition. At least 3 images per well were taken.

### Fluorescence microscopy and analysis

Cells were analyzed and imaged using a ZEISS Axio Observer Life Science Inverted Microscope while cells maintained at 38°C and 5% CO_2_. Images were analyzed and processed with Zen microscopy software 3.6 (ZEISS).

### Triglycerides quantification

Cellular triglyceride content was measured using a calorimetric assay (Cayman, 10010303-96). Triglyceride level for each well was normalized to protein concentrations via subsequent BCA (Bicinchoninic Acid) protein assay.

### Cell viability analysis

Cell viability was measured using CellTiter-Glo assay (Promega, G7573) according to the manufacturer recommendations. Briefly, cells were incubated with the reagent for 10 mins at 37 °C, and the luminescent signal was quantified by Infinite M Plex plate reader (Tecan).

### Statistical Analysis

Statistical analyses were performed using GraphPad Prism version 8.0.1. The specific test is described in the figure legend of each figure. A value of P < 0.05 was considered statistically significant.

Principal component analysis (PCA) scores plots were generated using MetaboAnalyst software 5.0 (https://www.metaboanalyst.ca/) with the annotated metabolomics data containing 270 compounds in the plasma and 350 in the liver from all samples. PCA plots were generated using log-transformed intensities and auto-scaling for normalizatio. Metabolite enrichment analysis was performed exclusively on annotated metabolites (HMDB ID) with statistically significant change (p-value < 0.05) using the Small Molecule Pathway Database (SMPDB) library and enrichment pathway tool in MetaboAnalyst software 5.0.

