## Supplemental figures and tables for "Fatty liver-mediated glycine restriction impairs glutathione synthesis and causes hypersensitization to acetaminophen"

### Supplementary Information

Supplementray figure.1:

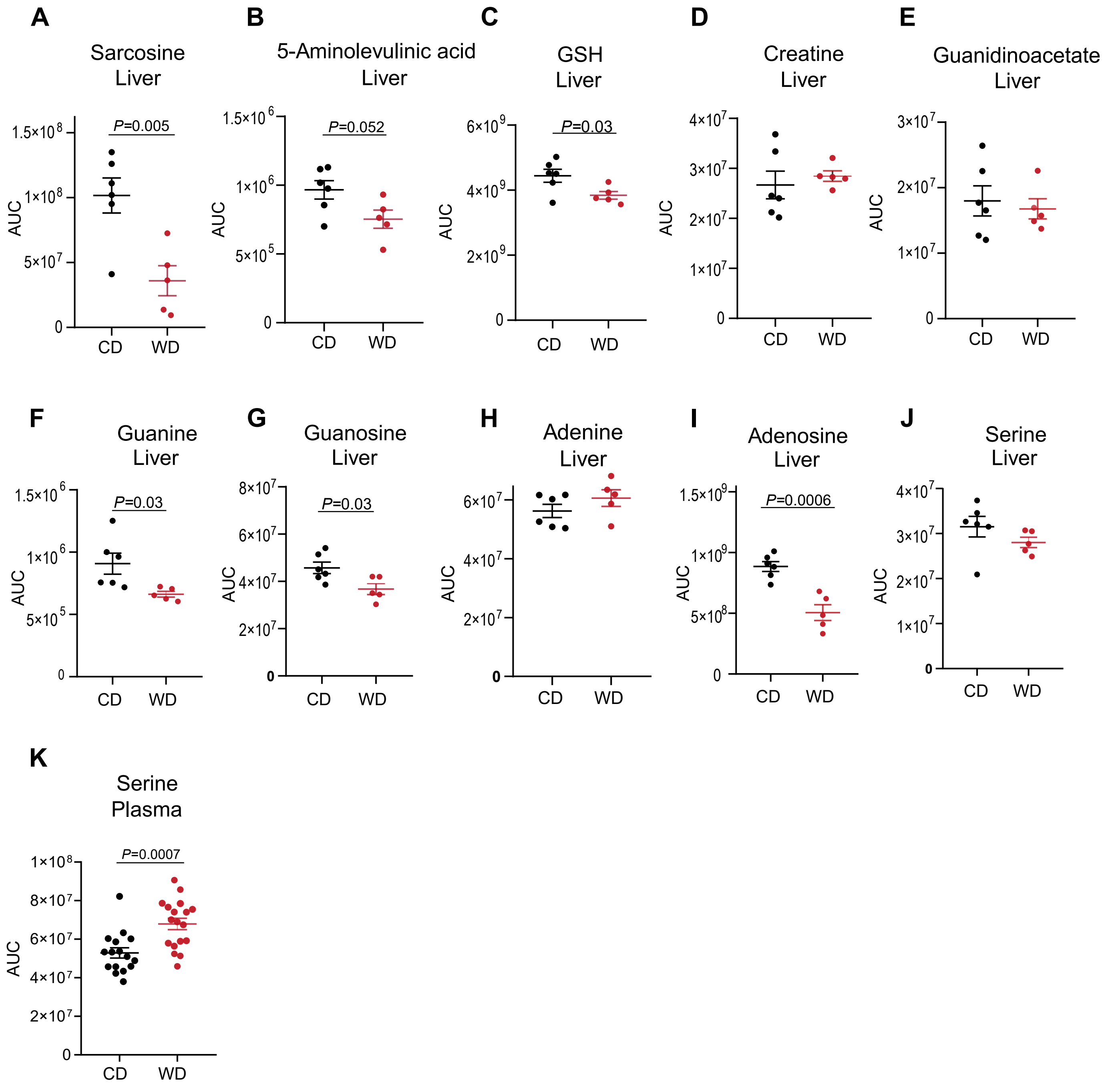

Supplementray figure.2:

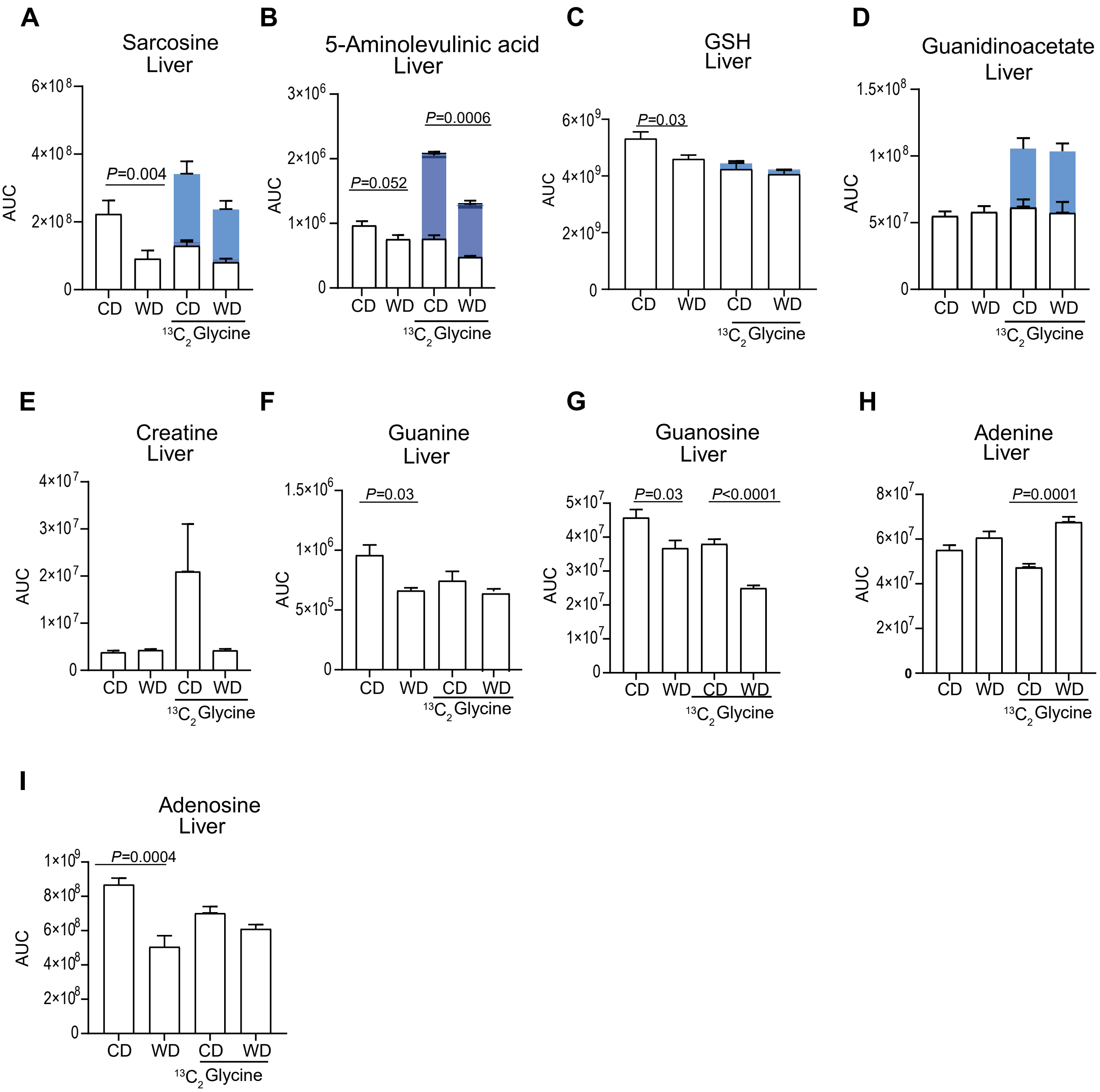

Supplementray figure.3:

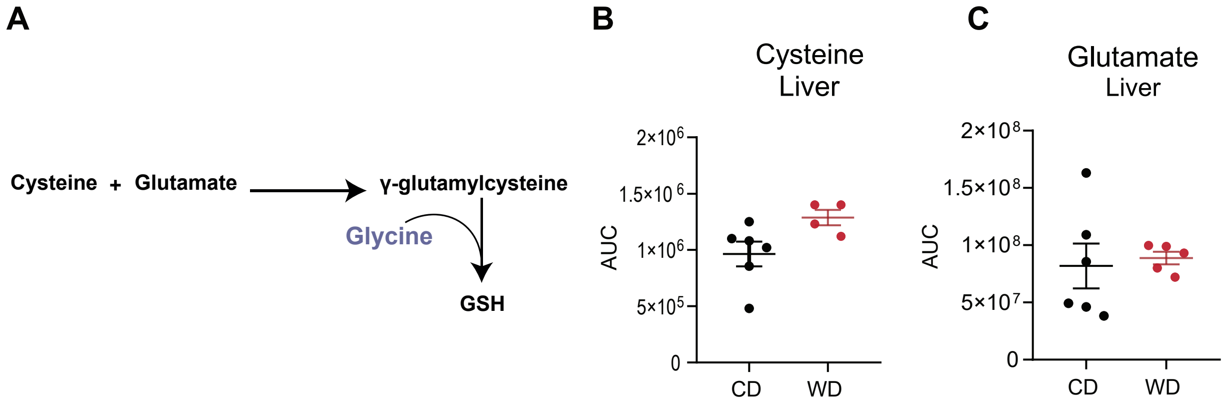

Supplementray figure.4:

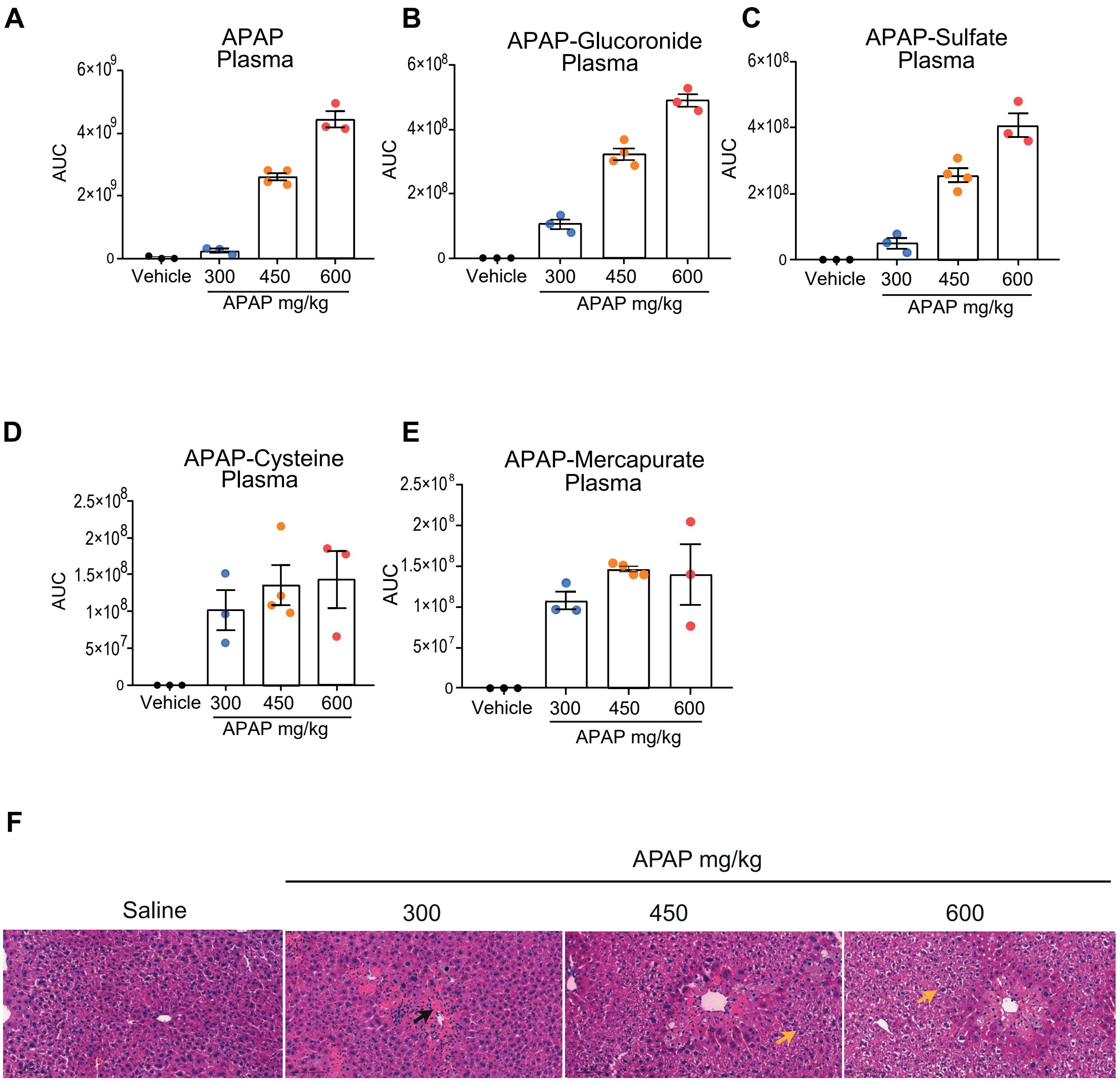

Supplementray figure.5:

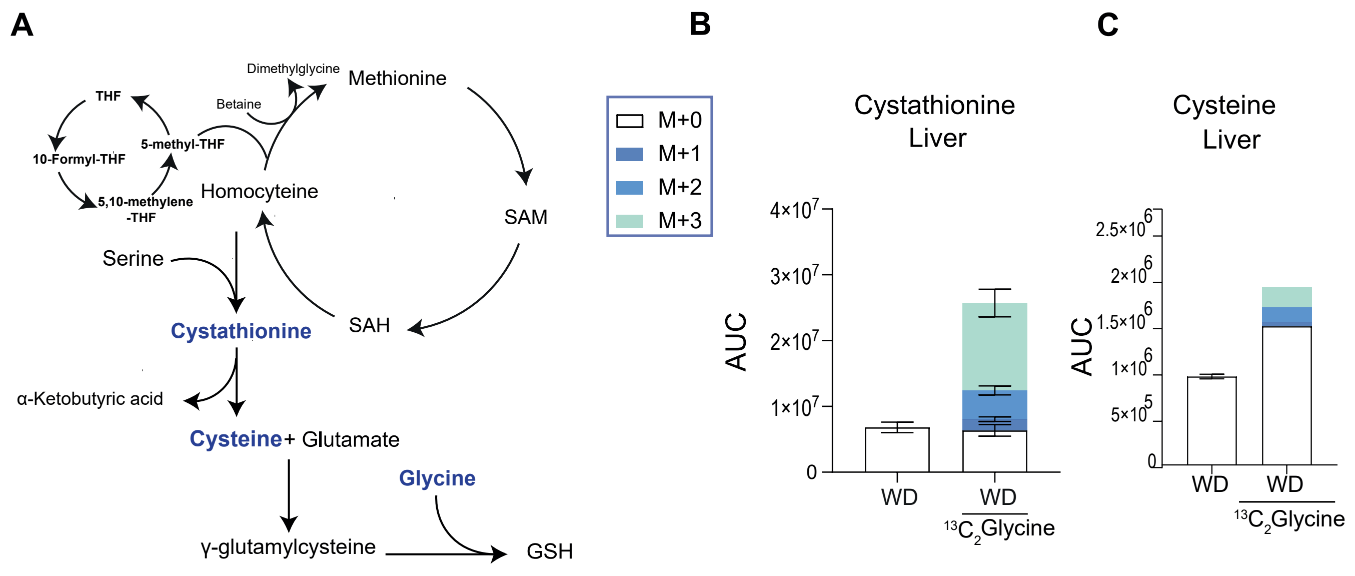

**Supplementary Figure Legend:**

**Figure S1:** (Related to Figure 1). **A-J,** Hepatic levels of glycine derived metabolites in CD and WD fed mice (n=6 and 5, respectively). **K,** Plasma levels of serine (n=16 in CD and n= 18 in WD group). Data are presented as mean ± SEM. Each point represents an individual mouse. *P* values were determined by two-tailed Student's *t*-test.

**Figure S2:** Mice were injected with $13C_2$ glycine and sacrificed 10 minutes following the 3rd injection of glycine (Related to Figure 3). **A-I,** Metabolic analysis of the liver samples (n= 6 in CD, n= 5 in WD, n=5 in CD-glycine and n=4 mice in WD-glycine group). Data are presented as mean ± SEM. Each point represents an individual mouse. *P* values were determined by ANOVA followed by Tukey's test.

**Figure S3** (Related to Figure 4). **A,** Schematic representation of GSH biosynthesis pathway. **B and C,** Hepatic levels of cysteine and glutamate in CD compared to WD fed mice (n= 6 and 5 respectively). Data are presented as mean ± SEM. Each point represents an individual mouse. *P* values were determined by two-tailed Student's *t*-test.

**Figure S4:** (Related to Figure 5). **A-E,** LC-MS analysis of APAP metabolic byproducts in the plasma (see Figure 4G) following 3 hrs of APAP administration at the indicated concentration (n= 3 per group). **F,** H&E-stained liver sections 24 hrs following APAP injection (scale bars: 100 μm, black arrow: necrotic area, yellow arrows: ballooning degeneration). Data are presented as mean ± SEM. Each point represents an individual mouse.

**Figure S5: A,** Schematic representation of transsulfuration pathway. **B and C,** Metabolic analysis of the livers following 3 injections of 500 mg/kg of $13C_2$ glycine. **B,** Cystathionine levels and C13 enrichment following 10 minutes of glycine injection (n=5 in CD and WD group, n=2 in CD+Glycine and n=3 in WD+Glycine group). **C,** Cysteine levels measured 3 hrs following glycine injection (n=5 in CD and WD group, n=1 in CD+Glycine and WD+Glycine group).

#### Supplementary Tables:

**Table S1:** (Related to Figure 2). List of detected metabolites in the liver with significant changes ($log_2$ fold change > 0.4 or < -0.4) in WD compared to CD fed mice (metabolites in red are methylated metabolites).

| Compound Name | Formula | Calculated molecular weight | RT [min] | Log2 fold change: | P-value |
|---|---|---|---|---|---|
| N-Methyl-2-pyrrolidone | C5 H9 N O | 99.06848 | 6.995 | -5.7 | < 0.0001 |
| Pyrogallol-2-O-glucuronide | C12 H14 O9 | 302.0636 | 2.8 | -5.3 | 0.0008 |
| Stachydrine | C7 H13 N O2 | 143.0947 | 3.374 | -4.7 | < 0.0001 |
| 1-Methylnicotinate | C7 H7 N O2 | 137.0477 | 4.368 | -4.5 | < 0.0001 |
| 4-Guanidinobutanoate | C5 H11 N3 O2 | 145.0854 | 9.135 | -4.3 | 0.0023 |
| N-Acetylornithine | C7 H14 N2 O3 | 174.1005 | 7.254 | -3.9 | < 0.0001 |
| L-Histidine trimethylbetaine | C9 H15 N3 O2 | 197.1165 | 5.091 | -3.8 | 0.0001 |
| Pipecolate | C6 H11 N O2 | 129.079 | 5.178 | -2.5 | 0.0001 |
| Propionyl-carnitine | C10 H19 N O4 | 217.1313 | 2.841 | -2.4 | 0.0286 |
| Hypotaurocyamine | C3 H9 N3 O2 S | 151.0416 | 9.579 | -2.1 | 0.0153 |
| Glycocholate | C26 H43 N O6 | 465.3099 | 1.259 | -2.0 | 0.0203 |
| Hypotaurine | C2 H7 N O2 S | 109.0198 | 8.851 | -1.9 | 0.0054 |
| Methyl 4-Aminobutyrate | C5 H11 N O2 | 117.0789 | 6.992 | -1.8 | 0.0085 |
| Eicosapentaenoate | C20 H30 O2 | 302.225 | 1.028 | -1.8 | < 0.0001 |
| 2-Aminoadipate | C6 H11 N O4 | 161.0689 | 9.887 | -1.8 | 0.0096 |
| Hydroxyhexanoycarnitine | C13 H25 N O5 | 275.1734 | 2.994 | -1.7 | 0.0009 |
| Acetyl-L-carnitine | C9 H17 N O4 | 203.1157 | 3.634 | -1.6 | 0.0233 |
| Butyryl-L-carnitine | C11 H21 N O4 | 231.147 | 2.432 | -1.6 | 0.0130 |
| Deoxycarnitine | C7 H15 N O2 | 145.1104 | 6.589 | -1.4 | < 0.0001 |
| 3-Hydroxyisovalerylcarnitine | C12 H23 N O5 | 261.1575 | 3.316 | -1.3 | 0.0018 |
| N-Acetylasparagine | C6 H10 N2 O4 | 174.0642 | 4.069 | -1.3 | 0.0056 |
| 3-hydroxybutyrylcarnitine | C11 H21 N O5 | 247.1419 | 4 | -1.3 | 0.0007 |
| Xanthosine | C10 H12 N4 O6 | 284.0757 | 6.052 | -1.3 | 0.0498 |
| Sarcosine | C3 H7 N O2 | 89.04775 | 7.619 | -1.3 | 0.04 |
| Adrenic acid | C22 H36 O2 | 332.2723 | 0.994 | -1.2 | 0.0010 |
| 6-Methylnicotinamide | C7 H8 N2 O | 136.0637 | 14.567 | -1.2 | 0.0221 |
| Methylarginine | C7 H16 N4 O2 | 188.1274 | 14.373 | -1.2 | < 0.0001 |
| Imidazolelactic acid | C6 H8 N2 O3 | 156.0537 | 3.808 | -1.2 | 0.0012 |
| Taurohyocholic acid | C26 H45 N O7 S | 515.2917 | 1.218 | -1.0 | 0.0207 |
| 5-guanidino-3-methyl-2-oxopentanoic acid | C7 H13 N3 O3 | 187.0958 | 7.786 | -1.0 | 0.0171 |
| N-Formylaspartate | C5 H7 N O5 | 161.0327 | 2.341 | -1.0 | 0.0119 |
| N-Alpha-Acetyllysine | C8 H16 N2 O3 | 188.1162 | 9 | -1.0 | 0.0005 |
| Pterin | C6 H5 N5 O | 163.0498 | 3.664 | -0.9 | 0.0428 |
| Dimethylglycine | C4 H9 N O2 | 103.0634 | 4.798 | -0.9 | 0.0191 |

| | | | | | |
|---|---|---|---|---|---|
| Imidazoleacetic acid | C5 H6 N2 O2 | 126.043 | 4.449 | -0.8 | 0.0007 |
| Cytidine | C9 H13 N3 O5 | 243.0856 | 4.814 | -0.8 | 0.0061 |
| Glycerol 3-Phosphate | C3 H9 O6 P | 172.0137 | 10.291 | -0.8 | 0.0012 |
| N-Acetyl-L-histidine | C8 H11 N3 O3 | 197.0802 | 1.405 | -0.8 | 0.0031 |
| Stearic acid | C18 H36 O2 | 284.2722 | 1.001 | -0.7 | 0.0512 |
| Acetylglycine | C4 H7 N O3 | 117.0426 | 2.967 | -0.7 | 0.0182 |
| N-Acetylneuraminate | C11 H19 N O9 | 309.1063 | 7.787 | -0.7 | 0.0090 |
| N-Acetylglucosaminitol | C8 H17 N O6 | 223.1058 | 6.334 | -0.7 | 0.0003 |
| Betaine | C5 H11 N O2 | 117.0789 | 3.866 | -0.7 | 0.0286 |
| Carnitine | C7 H15 N O3 | 161.1052 | 6.33 | -0.7 | 0.0003 |
| Methyl Vanillate | C9 H10 O4 | 182.0582 | 2.728 | -0.7 | 0.0054 |
| N(6)-Methyllysine | C7 H16 N2 O2 | 160.1213 | 14.301 | -0.7 | 0.0007 |
| Beta-Alanine | C3 H7 N O2 | 89.04777 | 9.361 | -0.7 | 0.0135 |
| Adenosine | C10 H13 N5 O4 | 267.0965 | 2.677 | -0.6 | 0.0127 |
| 3-Hydroxymethylglutarate | C6 H10 O5 | 162.0531 | 8.039 | -0.6 | 0.0191 |
| N-Acetylglutamate | C7 H11 N O5 | 189.0639 | 9.701 | -0.6 | 0.0311 |
| Glycine | C2 H5 N O2 | 75.03214 | 9.62 | -0.58 | 0.0002 |
| N-Acetylleucine | C8 H15 N O3 | 173.1055 | 1.279 | -0.5 | 0.0196 |
| Dihydrobiopterin | C9 H13 N5 O3 | 239.1018 | 5.427 | -0.5 | 0.0004 |
| 6-Methylpicolinate | C7 H7 N O2 | 137.0477 | 10.705 | -0.4 | 0.0039 |
| Methylimidazoleacetic acid | C6 H8 N2 O2 | 140.0586 | 2.613 | -0.4 | 0.0074 |
| 2-Methyl-beta-alanine | C4 H9 N O2 | 103.0634 | 7.077 | -0.4 | 0.0365 |
| Dihydrothymine | C5 H8 N2 O2 | 128.0586 | 8.758 | -0.4 | 0.0076 |
| Dimethylarginine | C8 H18 N4 O2 | 202.1431 | 13.286 | -0.4 | 0.0034 |
| N6-Acetyl-L-lysine | C8 H16 N2 O3 | 188.1161 | 6.131 | -0.4 | 0.0337 |
| Fructose 6-Phosphate | C6 H13 O9 P | 260.0299 | 11.205 | 0.35 | 0.0055 |
| Aspartate | C4 H7 N O4 | 133.0375 | 9.911 | 0.36 | 0.0754 |
| Galacturonate | C6 H10 O7 | 194.043 | 10.363 | 0.39 | 0.0280 |
| Uridine 5'-Diphosphate | C9 H14 N2 O12 P2 | 404.002 | 11.564 | 0.41 | 0.0228 |
| Deoxycholate | C24 H40 O4 | 392.2931 | 2.294 | 0.42 | 0.0001 |
| Adenine | C5 H5 N5 | 135.0545 | 2.823 | 0.42 | 0.0106 |
| Saccharate | C6 H10 O8 | 210.0379 | 11.745 | 0.44 | 0.0320 |
| Citrate | C6 H8 O7 | 192.0269 | 12.65 | 0.44 | 0.0194 |
| Uridine Diphosphate Glucose | C15 H24 N2 O17 P2 | 566.0556 | 11.564 | 0.49 | 0.0084 |
| Succinylaminoimidazole carboxamide riboside | C13 H18 N4 O9 | 374.1074 | 11.892 | 0.5 | 0.0263 |
| L-Methionine sulfoxide | C5 H11 N O3 S | 165.0461 | 6.953 | 0.5 | 0.0545 |
| Mesaconic acid | C5 H6 O4 | 130.0268 | 10.586 | 0.53 | 0.0034 |
| Palmitamide | C16 H33 N O | 255.2563 | 24.155 | 0.53 | 0.0067 |
| L-Glutamic acid | C5 H9 N O4 | 147.0535 | 12.08 | 0.54 | 0.0067 |
| 2-hydroxy glutarate | C5 H8 O5 | 148.0376 | 10.59 | 0.54 | 0.0235 |
| Sulfuric acid | H2 O4 S | 97.96773 | 12.404 | 0.55 | 0.0275 |
| DL-gamma-carboxyglutamic acid | C6 H9 N O6 | 191.0431 | 12.078 | 0.61 | 0.0020 |
| Leucine | C6 H13 N O2 | 131.0946 | 3.659 | 0.63 | 0.0158 |
| Homocitrulline | C7 H15 N3 O3 | 189.1114 | 8.865 | 0.65 | 0.0076 |
| Guanidinoethyl sulfonate | C3 H9 N3 O3 S | 167.0365 | 9.52 | 0.71 | 0.0057 |

| | | | | | |
|---|---|---|---|---|---|
| Hydroxycitric acid | C6 H8 O8 | 208.0221 | 11.306 | 0.73 | 0.0159 |
| O-Phosphoethanolamine | C2 H8 N O4 P | 141.0192 | 10.839 | 0.74 | 0.0023 |
| S-Adenosylmethionine | C15 H22 N6 O5 S | 398.1371 | 10.804 | 0.74 | 0.0041 |
| Galactarate | C6 H10 O8 | 210.038 | 12.087 | 0.79 | 0.0062 |
| L-Threonic acid | C4 H8 O5 | 136.0372 | 6.144 | 0.99 | 0.0181 |
| Methylthioadenosine | C11 H15 N5 O3 S | 297.0893 | 2.338 | 1.01 | 0.0024 |
| N-[(2S)-2-Hydroxypropanoyl]methionine | C8 H15 N O4 S | 221.0721 | 9.766 | 1.09 | 0.0053 |
| Myristic acid | C14 H28 O2 | 228.2093 | 1.063 | 1.41 | 0.0378 |
| Glycerophosphoglycerol | C6 H15 O8 P | 246.0506 | 6.92 | 1.55 | 0.0001 |
| Cholesterol sulfate | C27 H46 O4 S | 466.3118 | 0.926 | 1.62 | < 0.0001 |
| Acetylaspartic acid | C6 H9 N O5 | 175.0483 | 10.083 | 1.63 | < 0.0001 |
| Arginine | C6 H14 N4 O2 | 174.1118 | 15.485 | 1.79 | 0.0038 |
| Lysophosphatidic acid | C21 H41 O7 P | 436.2589 | 2.288 | 1.82 | 0.0007 |
| Eicosenoic acid | C20 H38 O2 | 310.2877 | 0.985 | 1.85 | 0.0065 |
| Oleic acid | C18 H34 O2 | 282.256 | 1.015 | 1.93 | 0.0179 |
| Palmitoleic acid | C16 H30 O2 | 254.225 | 1.049 | 2.15 | 0.0165 |
| trans-10-Heptadecenoic Acid | C17 H32 O2 | 268.2405 | 1.034 | 2.31 | 0.0118 |
| 1-[(9Z)-hexadecenoyl]-sn-glycero-3-phosphocholine | C24 H48 N O7 P | 493.3169 | 2.299 | 2.34 | 0.0002 |
| Ureidopropionate | C4 H8 N2 O3 | 132.0535 | 5.336 | 2.49 | 0.0058 |
| Sphinganine | C18 H39 N O2 | 301.2978 | 2.302 | 2.5 | < 0.0001 |
| Ascorbic acid | C6 H8 O6 | 176.0323 | 12.864 | 3.86 | < 0.0001 |

**Table S2:** (Related to Figure 2). List of detected metabolites in the plasma with significant changes ($log_2$ fold change > 0.4 or < -0.4) in WD compared to CD fed mice (metabolites in red are methylated metabolites).

| Compound Name | Formula | Calculated molecular weight | RT [min] | Log2 Fold Change | P-value |
|---|---|---|---|---|---|
| Hypoxanthine | C5 H4 N4 O | 136.0386 | 3.352 | -9.03 | < 0.0001 |
| Inosine | C10 H12 N4 O5 | 268.0807 | 4.054 | -8.48 | < 0.0001 |
| Xanthine | C5 H4 N4 O2 | 152.0339 | 4.255 | -8.45 | < 0.0001 |
| 1-Methylnicotinate | C7 H7 N O2 | 137.0478 | 4.348 | -5.28 | < 0.0001 |
| N-Acetylornithine | C7 H14 N2 O3 | 174.1005 | 7.373 | -4.64 | < 0.0001 |
| Dihydrobiopterin | C9 H13 N5 O3 | 239.1018 | 3.403 | -3.84 | < 0.0001 |
| Indole-3-Methyl Acetate | C11 H11 N O2 | 189.0793 | 2.349 | -3.3 | 0.0148 |
| Xanthosine | C10 H12 N4 O6 | 284.0757 | 6.704 | -3.11 | < 0.0001 |
| Linolenic acid | C18 H30 O2 | 278.2249 | 1.037 | -3.1 | < 0.0001 |
| Tyramine glucuronide | C14 H19 N O7 | 313.116 | 10.603 | -2.71 | < 0.0001 |
| Linoleic acid | C18 H32 O2 | 280.2404 | 1.025 | -2.71 | < 0.0001 |
| Acetyl-β-methyl choline | C8 H17 N O2 | 159.1262 | 6.513 | -2.15 | < 0.0001 |
| N-Acetyl-L-glutamine | C5 H5 N5 | 135.0546 | 2.844 | -2.12 | 0.0003 |
| g-Butyrobetaine | C7 H15 N O2 | 145.1104 | 2.847 | -2.04 | < 0.0001 |
| Docosahexaenoic Acid | C22 H32 O2 | 328.2407 | 1.01 | -1.46 | < 0.0001 |
| Methylimidazole acetic acid | C6 H8 N2 O2 | 140.0587 | 3.415 | -1.25 | 0.0002 |
| N-Methyl glutamate | C6 H11 N O4 | 161.069 | 6.966 | -1.22 | < 0.0001 |
| Arachidonic acid | C20 H32 O2 | 304.2406 | 1.022 | -1.2 | < 0.0001 |
| Glycerophosphoethanolamine | C23 H44 N O7 P | 477.2859 | 2.306 | -1.13 | 0.0002 |
| N-Alpha-Acetyl lysine | C8 H16 N2 O3 | 188.1162 | 8.999 | -1.1 | < 0.0001 |
| N-Acetyl leucine | C8 H15 N O3 | 173.1055 | 1.272 | -1.03 | 0.0183 |
| Propionyl-carnitine | C10 H19 N O4 | 217.1314 | 2.85 | -1 | < 0.0001 |
| N-Acetyl glycine | C4 H7 N O3 | 117.0427 | 3.052 | -0.87 | 0.0008 |
| Citrate | C6 H8 O7 | 192.0273 | 12.591 | -0.82 | 0.0001 |
| Glutaurine | C7 H14 N2 O6 S | 254.0574 | 10.329 | -0.82 | 0.0003 |
| Homocitrulline | C7 H15 N3 O3 | 189.1114 | 9.339 | -0.82 | 0.0001 |
| 3-Hydroxybutanoate | C4 H8 O3 | 104.0477 | 2.568 | -0.77 | 0.0119 |
| N-Methyl glutamine | C6 H12 N2 O3 | 160.0849 | 6.853 | -0.76 | 0.0004 |
| Acetyl-L-carnitine | C9 H17 N O4 | 203.1157 | 3.657 | -0.68 | < 0.0001 |
| N (6)-Methyl lysine | C7 H16 N2 O2 | 160.1213 | 14.149 | -0.68 | 0.0073 |
| Deoxy carnitine | C7 H15 N O2 | 145.1104 | 6.575 | -0.67 | < 0.0001 |
| Methylarginine | C7 H16 N4 O2 | 188.1275 | 14.157 | -0.64 | 0.0013 |
| Hydroxybutyrylcarnitine | C11 H21 N O5 | 247.1419 | 3.971 | -0.61 | 0.0023 |
| Acetyl arginine | C8 H16 N4 O3 | 216.1222 | 8.898 | -0.56 | < 0.0001 |
| Pipecolate | C6 H11 N O2 | 129.079 | 5.162 | -0.53 | 0.0002 |
| Malate | C4 H6 O5 | 134.0215 | 11.098 | -0.52 | 0.0013 |

| | | | | | |
|---|---|---|---|---|---|
| Carnitine | C7 H15 N O3 | 161.1052 | 6.333 | -0.5 | < 0.0001 |
| N,N-Dimethyl lysine | C8 H18 N2 O2 | 174.137 | 12.727 | -0.46 | 0.0128 |
| Betaine | C5 H11 N O2 | 117.0789 | 3.862 | -0.43 | 0.0016 |
| Cis-Aconitate | C6 H6 O6 | 174.0166 | 12.185 | -0.43 | < 0.0001 |
| Glycine | C2 H5 N O2 | 75.03221 | 9.598 | -0.43 | 0.0001 |
| Palmitate | C16 H32 O2 | 256.2406 | 1.053 | -0.43 | 0.0038 |
| Methylbutyroyl-carnitine | C12 H23 N O5 | 261.1575 | 3.319 | -0.42 | 0.0208 |
| Methionine | C5 H11 N O2 S | 149.0512 | 4.224 | 0.43 | 0.03 |
| 6-Methylquinoline | C10 H9 N | 143.0736 | 4.587 | 0.43 | 0.0247 |
| Kynurenine | C10 H12 N2 O3 | 208.0848 | 3.825 | 0.57 | 0.0234 |
| Trans-10-Heptadecenoic Acid | C17 H32 O2 | 268.2405 | 1.033 | 0.94 | 0.0005 |
| L-alpha-Lys phosphatidylcholine | C22 H46 N O7 P | 467.3012 | 2.334 | 2.04 | < 0.0001 |
| Dimethylguanidino valeric acid | C8 H15 N3 O3 | 201.1114 | 3.885 | 0.4 | 0.0212 |
| N-Acetyl tryptophan | C13 H14 N2 O3 | 246.1005 | 2.352 | 0.57 | 0.0029 |
| Glucose | C6 H12 O6 | 180.0636 | 8.039 | 0.45 | 0.0038 |
| Pentadecanoic acid | C15 H30 O2 | 242.2249 | 1.056 | 0.63 | 0.0044 |
| Asparagine | C4 H8 N2 O3 | 132.0536 | 9.318 | 0.59 | 0.0080 |
| Ascorbic acid | C6 H8 O6 | 176.0323 | 12.735 | 1.84 | 0.0026 |
| Phenylacetyl glycine | C10 H11 N O3 | 193.0743 | 2.341 | 0.81 | 0.0362 |
| Myristoleic acid | C14 H26 O2 | 226.1935 | 1.066 | 1.59 | < 0.0001 |
| Serine | C3 H7 N O3 | 105.0427 | 9.652 | 0.41 | 0.0004 |
| Ornithine | C5 H12 N2 O2 | 132.0899 | 13.967 | 0.47 | 0.1198 |
| Decanoic acid | C10 H20 O2 | 172.1466 | 1.113 | 0.84 | 0.0015 |
| L-Threonic acid | C4 H8 O5 | 136.0372 | 6.749 | 0.53 | 0.0285 |
| 3-Methyl-2-Oxovalerate & Ketoleucine | C6 H10 O3 | 130.0632 | 1.285 | 0.46 | 0.0525 |
| Lauric acid | C12 H24 O2 | 200.178 | 1.082 | 2.1 | < 0.0001 |
| Butyryl-L-carnitine | C11 H21 N O4 | 231.147 | 2.457 | 1.09 | < 0.0001 |
| Myristic acid | C14 H28 O2 | 228.2092 | 1.064 | 0.82 | 0.0013 |
| Pantothenate | C9 H17 N O5 | 219.1107 | 2.38 | 0.55 | 0.0405 |
| Trans-3-Indoleacrylic acid | C11 H9 N O2 | 187.0634 | 4.594 | 0.43 | 0.0244 |
| Threonine | C4 H9 N O3 | 119.0582 | 8.095 | 0.57 | 0.0073 |
| Indole-3-Pyruvate | C11 H9 N O3 | 203.0587 | 2.348 | 0.75 | 0.0009 |

**Supplementary movies:**

**Supplementary movie. 1 .**
